# Proteome-wide Mapping of Cysteine Persulfidation in Deep-Sea Hyperthermophilic Archaea

**DOI:** 10.64898/2026.01.05.697748

**Authors:** Ling Fu, Xuegong Li, Shaowei Liu, Yunhan Zhang, Jing Yang

## Abstract

Protein persulfidation, a primary post-translational modification mediated by the gaseous signaling molecule hydrogen sulfide (H_2_S), regulates diverse physiological processes in eukaryotes and bacteria. However, its existence and functional roles in archaea, the third domain of life, remain completely unexplored. Here, we investigate this in the deep-sea hyperthermophilic archaeon *Thermococcus aciditolerans* SY113, which thrives in sulfur rich hydrothermal vents and endogenously produces substantial H_2_S. Our profiling not only delineated the unique reactivity landscape of these modifications but also enabled quantitative analysis of its dynamic regulation by H_2_S. A total of 204 persulfidation sites on 171 proteins were identified, over 65% of which were dynamically regulated by H_2_S. Further functional analysis suggests that persulfidation represents an ancient and conserved regulatory mechanism in primordial life including the regulation of catalytic activity, maintenance of protein conformation, and mediation of protein-protein interactions. Our findings provide a valuable dataset and theoretical foundation for understanding the role of persulfidation in the physiological regulation of deep-sea hyperthermophilic archaea.

## Introduction

Deep-sea hydrothermal vents are among Earth’s most extreme and unique ecosystems, characterized by perpetual darkness, high pressure, and steep physicochemical gradients[1, 2]. Despite these challenging conditions, vents support thriving biological communities largely dependent on chemolithoautotrophic microorganisms that utilize inorganic chemicals as energy sources[3]. These microbes oxidize reduced inorganic compounds such as hydrogen sulfide (H_2_S) and hydrogen (H_2_) released from the vents, to gain energy and synthesize organic carbon, thereby acting as primary producers and forming the foundation of the hydrothermal vent food web[4]. Consequently, chemolithoautotrophs play an indispensable role in material and energy transformation, biogeochemical cycling, and the maintenance of the vent ecosystem[5].

Within this context, archaea of the order Thermococcales constitute a major group of deep-sea hyperthermophilic chemolithoautotrophs [6, 7]. These organisms preferentially use elemental sulfur (S⁰) as a terminal electron acceptor, with their growth being significantly stimulated by its presence [8, 9]. This metabolic process is tightly regulated by the transcription factor SurR[10, 11]. Under reducing conditions, SurR binds to promoter elements containing a “GTTn_₃_AAC” motif, repressing sulfur metabolism genes. In the presence of S⁰, however, redox-sensitive cysteine residues within its “CXXC” motif undergo conformational changes, leading to derepression and upregulation of key sulfur metabolic enzymes, including membrane-bound sulfane reductase (MBS) and NADPH sulfur oxidoreductase (NSR)[12, 13]. These enzymes function synergistically: MBS uses reduced ferredoxin to generate NADPH, which NSR then employs to reduce S⁰, producing substantial amounts of H_2_S [14, 15]. This regulatory pathway clearly indicates that H_2_S is not only a key metabolic by-product but also a potential signaling molecule involved in energy metabolism and growth regulation in Thermococcales.

H_2_S is recognized as the third gaseous signaling molecule, following nitric oxide (NO) and carbon monoxide (CO), and is considered an important component in early life evolution[16]. In eukaryotic systems, H_2_S regulates diverse physiological processes through a primary molecular mechanism known as persulfidation, wherein H_2_S modifies reactive cysteine thiols to form persulfide groups (-SSH)[17–19]. This modification can alter protein function, stability, localization, and interaction networks, underpinning the broad regulatory roles of H_2_S[20]. To date, research on H_2_S signaling and persulfidation, has been largely confined to eukaryotes and a limited number of bacteria. Thus, a fundamental question arises: in these prolific H_2_S-producing archaea, does persulfidation, serve as an intrinsic signaling mechanism, analogous to its role in eukaryotes? Addressing this question is crucial for understanding the evolution of redox signaling and the adaptive strategies of life in extreme environments.

We therefore hypothesized that persulfidation constitutes a fundamental signaling mechanism in Thermococcales. To test this, we employed the hyperthermophilic archaeon *Thermococcus aciditolerans* SY113, which isolated from the Longqi hydrothermal field in the Southwest Indian Ocean at a depth of 2770 meters, and applied chemical proteomics to systematically profile protein persulfidation in archaea for the first time[21]. Our study not only confirms the widespread presence of this post-translational modification in archaea but also reveals unique functional features distinct from those observed in eukaryotes and bacteria. We further demonstrate that persulfidation modulates enzymatic activity, maintains protein conformation, and mediates protein-protein interactions, suggesting an ancient and essential regulatory role in primordial life forms[22].

## Materials and methods

### Strains and growth conditions

*T. aciditolerans* SY113 was grown at 85_℃_ in TRM liquid medium under anaerobic conditions[21]. To examine the growth characteristics of the SY113 strain, the culture was cultivated in TRM medium supplemented with elemental sulfur (S⁰) until it reached the stationary phase (OD_600nm_≈0.6). The culture was then centrifuged at 2,000 × g and 4 °C for 10 min to remove residual sulfur powder. The supernatant was transferred to a new tube and subjected to a second centrifugation at 5,000 × g and 4 °C for 15 min. The resulting cell pellet was collected and washed three times with fresh anoxic TRM medium inside an anaerobic chamber. Subsequently, the cells were resuspended in anaerobic TRM medium to an optical density at 600 nm (OD_₆₀₀_) of approximately 1.0. This cell suspension was used as an inoculum at 1% (v/v) in fresh TRM medium. For sulfur-supplemented conditions, elemental sulfur was added to a final concentration of 2 g/L. High-pressure cultivation at 27 MPa was conducted using a custom stainless-steel pressure vessel (Nantong Feiyu Petroleum Technology Development Co., Ltd., China) equipped with a pressure gauge and an automated water-pumping system for precise pressure control.

### Probe labeling and proteomic sample preparation

For analyzing intrinsic SSH-reactivity, SY113 pellets were lysed by sonication in four volumes of lysis buffer (50 mM HEPES pH 4.5, 150 mM NaCl, and 1% Igepal) supplemented with 1x protease and phosphatase inhibitors and 200 unit/mL catalase. Cell lysates were divided into two parts and incubated with 10 μM or 100 μM IPM at RT for 1 hr with rotation and light protection, respectively.

For analyzing SSH dynamics, SY113 pellets with or without sulfur addition were lysed by sonication in four volumes of lysis buffer (50 mM HEPES pH 4.5, 150 mM NaCl, and 1% Igepal) supplemented with 1x protease and phosphatase inhibitors and 200 unit/mL catalase. Cell lysates were then lebeled with 100 μM IPM at RT for 1 hr with rotation and light protection, respectively.

After incubation, samples were alkylated with 40 mM IAA (Sigma-Aldrich, cat. no. I1149) for 30 min in the dark. The reactions were quenched by protein precipitation, which was performed with a methanol-chloroform system (aqueous phase/methanol/chloroform, 4:4:1, v/v/v). Proteins were collected via centrifugation at 1,700 x g for 20 min at 4°C. Liquid layers were discarded and the protein was washed twice in methanol/chloroform (1:1, v/v), followed by centrifugation at 12,000 x g for 10 min at 4 °C to repellet the protein. The protein pellets were resuspended with 50 mM ammonium bicarbonate (NH_4_HCO_3_, Sigma-Aldrich, A6141). Protein concentrations were measured with BCA assay kit (Tiangen, PA115) and adjusted to 2 mg/mL. Each of the control and antibiotic treated proteome samples (2 mg protein/mL in 0.5 mL volume, total: 1 mg each) were digested with sequencing grade trypsin (Sigma-Aldrich, A6141) at a 1:50 (enzyme/substrate) ratio for 16 h at 37 °C. The tryptic digests were evaporated to 200 uL and desalted with HLB extraction cartridges (Waters, cat. no. 186000383). The desalted tryptic digests were reconstituted in a solution containing 30 % ACN (Sigma-Aldrich, A6141). CuAAC reaction was performed by the addition of 0.8 mM either Azido-L-biotin or Azido-H-biotin, 8 mM sodium ascorbate (Sigma-Aldrich, A7631), 1 mM TBTA (Sigma-Aldrich, 678937), and 8 mM CuSO_4_ (Sigma-Aldrich, 678937). Samples were allowed to react at room temperature for 2 h in the dark with rotation. The light- and heavy Az-UV-biotin labeled samples then were mixed equally together. The excess biotin reagents were removed by strong cation exchange (SCX) spin columns (The Nest Group, SMM HIL-SCX). The sample was diluted into SCX loading buffer (5 mM KH_2_PO_4_, 25 % acetonitrile, pH 3.0), passed through the SCX spin columns, and washed with several column volumes of loading buffer. The retained peptides were eluted with SCX loading buffer containing 400 mM NaCl. Eluent was diluted 10 x with 50 mM sodium acetate buffer (NaAc, pH 4.5) and then allowed to interact with pre-washed streptavidin sepharose (Cytiva, cat. no. 17-5113-0) for 2 h at room temperature. Streptavidin sepharose then was washed with 50 mM NaAc (NaOAc, Sinopharm Chemical Reagent, cat. no. 10018818), 50 mM NaAc containing 2 M NaCl, and water (Fisher Scientific, cat. no. W5-4) twice each with vortexing and/or rotation to remove non-specific binding peptides, and resuspended in 25 mM fresh NH_4_HCO_3_. The suspension of streptavidin sepharose was transferred to several thin-walled borosilicate glass tubes, irradiated with 365 nm UV light (UVP, UVL-28 EL series) for 2h at room temperature with stirring. The supernatant was collected, concentrated under vacuum, and desalted with HLB extraction cartridges as previously described. The desalting peptides were evaporated to dryness and stored at -20 °C until LC-MS/MS analysis.

### LC-MS/MS

LC-MS/MS analyses were performed on Q Exactive plus (Thermo Fisher Scientific) operated with an Easy-nLC1000 system (ThermoFisher Scientific). Samples were reconstituted in 0.1 % formic acid (FA, Fisher Scientific, cat. no. A117-50) followed by centrifugation (16,000 g for 10 min) and the supernatants were pressure-loaded onto a 2 cm microcapillary precolumn packed with C18 (3 μm, 120 Å, SunChrom, USA). The precolumn was connected to a 12 cm 150-μm-inner diameter microcapillary analytical column packed with C18 (1.9 μm, 120 Å, Dr. MaischGebH, Germany) and equipped with a homemade electrospray emitter tip. The spray voltage was set to 2.0 kV and the heated capillary temperature to 320 °C. LC gradient consisted of 0 min, 7 % B; 14 min, 10 % B; 51 min, 20 % B; 68 min, 30 % B; 69-75 min, 95 % B (A = water, 0.1 % formic acid; B = ACN, 0.1 % formic acid) at a flow rate of 600 nL/min. HCD MS/MS spectra were recorded in the data dependent mode using a Top 20 method. MS1 spectra were measured with a resolution of 70,000, an AGC target of 3e6, a max injection time of 20 ms, and a mass range from m/z 300 to 1400. HCD MS/MS spectra were acquired with a resolution of 17,500, an AGC target of 1e6, a max injection time of 60 ms, a 1.6 m/z isolation window and normalized collision energy of 30. Peptide m/z that triggered MS/MS scans were dynamically excluded from further MS/MS scans for 18 s.

### Peptide Identification and Quantification

Raw data files were searched against the SY113 UniProt canonical database (20210528) using pFind studio (v3.1.5, http://pfind.ict.ac.cn/software/pFind/index.html). The precursor-ion mass and fragmentation tolerance were 10 ppm. and 20 ppm., respectively, for the database search. A specific-tryptic search was used with a maximum of three missed cleavages allowed. A maximum of three modifications were allowed per peptide. Modifications of +15.9949 Da (methionine oxidation, M) and +57.0214 Da (IAM alkylation, C) were searched as variable modifications. For site-specific mapping of IPM-modified sites, mass shifts of + 284.0942 (C_11_H_16_O_3_N_4_S_1_) for IPM was searched as variable modifications, respectively. A differential modification of 6.0201 Da on probe-derived modifications was used for all analyses. The FDRs at spectrum, peptide, and protein level were <1 %. Quantification of heavy to light ratios (R_H/L_) was performed using pQuant as previously described, which directly uses the RAW files as the input. pQuant calculated R_H/L_ values based on each identified MS scan with a 15 ppm-level m/z tolerance window and assigned an interference score (Int. Score) to each value from zero to one. The median values of probe-modified peptide ratios with σ less than or equal 0.5 were considered to calculate site-level ratios. For obtaining the site-centric quantification data, the output reports from pFind were then further processed using in-house software written in the R programming language as previously described[23].

### H_2_S detection

SY113 were grown to an OD_600_ of ∼0.5 and then pelleted at 12000 g for 5 min. H_2_S was determined with a commercialized assay kit (Solarbio, cat. no. BC2055) following the manufacturer’s instructions. The measurement of the absorption (665 nm) derived from a colorimetric reaction was conducted on a Multiskan MK3 plate reader.

### Construction of protein expression plasmids

To construct the protein expression plasmids pET28a (+)-*gpdh*, the full-length *gpdh* was amplified from SY113 genomic DNA respectively using corresponding primer pair. The DNA fragments were digested with indicated endonuclease and then cloned into the expression vector pET28a (+). Recombinant plasmid was introduced into the *E. coli* BE21(DE3) strain (TRANS, cat. no.CD601-02) to express the recombinant protein. The clones contained recombinant plasmids were cultured in 15 mL of Luria-Bertani (LB) medium at 37 °C and 200 rpm until OD_600_=1.5∼2. 5 mL of culture was inoculated into 500 mL of LB medium and further cultured at 37 °C and 200 rpm until OD_600_ = 0.6. After cooling at 30 °C for 20 min, isopropylthio-β-galactoside (0.5 mM final concentration, TRANS, cat. no.GF101-01) was added to the culture to induce the expression of recombinant proteins at 30 °C for 4 h. Recombinant proteins were purified via immobilized Ni^2+^ affinity chromatography. In brief, the bacterial pellet was resuspended in lysis buffer (20 mM Tris-HCl, pH 8.0, 300 mM NaCl, 5 mM β-mercaptoethanol, 10 mM imidazole, 1 mM phenylmethylsulfonyl fluoride, and 10 % glycerol (v/v)) and then disrupted by sonication. The cell extracts were clarified by centrifugation at 12,000 × g for 30 min at 4 °C. After loading the supernatant onto a Ni^2+^ affinity column pre-equilibrated with lysis buffer, the resin was washed with 6 column volumes of lysis buffer containing 20 mM imidazole. Next, the bound protein was eluted from the column using elution buffer (20 mM Tris-HCl, pH 8.0, 300 mM NaCl, 5 mM β-mercaptoethanol, 200 mM imidazole, and 10 % glycerol (v/v). The eluted protein was subsequently further purified by Superdex 200 Increase 10/300 GL (Cytiva, cat. no.28990944) gel filtration column chromatography. The purity of the elutes were verified by 12 % SDS-PAGE. Finally, the proteins were stored in small aliquots at -80 °C for activity detection.

### Enzyme activity assays

Recombinant protein was reduced with 1 mM DTT for 30 min at 4 °C. Reactions were then passed through Zeba™ Spin Desalting Column equilibrated with PBS, pH 7.4. DTT-free proteins were then treated with hydrogen peroxide (H_2_O_2_, 5 eq. Sigma-Aldrich, cat. no. 216763) at 37 _℃_ for 30 min. Subsequently, 0.1 mM of NaHS was added for 1hr at 37 _℃_. For GAPDH, the activity of GAPDH enzyme was quantified using a GAPDH activity assay kit (Abcam, cat. no. ab204732). Briefly, 2 μg GAPDH was used to measure GAPDH enzymatic activity using 96 well plates according to the manufacturer’s protocol. Absorbance was measured at 450 nm in kinetic mode with a microplate reader. GAPDH activity was calculated according to the manufacturer’s guidelines. Positive control provided by the kit manufacturer (to ensure the activity of kit components) was used as the control. For SAT, Enzymatic activity of SAT was assayed by molybdenum-dependent formation of phosphate. The reaction started by adding 15 mM of Na_2_MoO_4_ to the reaction mixture with 10 μg of SAT. The reaction mixture was consisted of PBS (pH7.4), Na_2_MoO_4_ (15 mM, Sigma-Aldrich, cat. no. 243655-5G), Na_2_ATP (2 mM, Sigma-Aldrich, cat. no. A26209), MgCl_2_ (7 mM), and inorganic pyrophosphatase (0.33 U × ml^−1^, Sigma-Aldrich, cat. no. I1643). The reaction was stopped after 15 min of incubation at 37°C by adding 1.0 ml of ice cold 0.5 mM sodium acetate (pH 4.0) and 200 μl of developer solution which consist of L-ascorbic acid (200 mg), Na_2_MoO_4_ (100 mg) in 10 ml of 0.36 M sulfuric acid (Sigma-Aldrich, cat. no. 339741). Absorbance was measured at 660 nm against blank PBS buffer.

### Transcriptome sequencing

SY113 was cultured in TRM medium with or without elemental sulfur (S⁰) supplementation until the stationary phase was reached. Total RNA was extracted using the RNAprep Pure Cell/Bacteria Kit (Tiangen, Beijing, China). RNA quality was verified by 1% agarose gel electrophoresis, while RNA concentration and integrity were evaluated with the RNA Nano 6000 Assay Kit on the Bioanalyzer 2100 system (Agilent Technologies, CA, USA). Ribosomal RNA was depleted prior to cDNA synthesis. After Illumina NovaSeq 6000 sequencing, the obtained cDNA sequences were aligned to the reference genome (see Genome sequencing section). Differential gene expression analysis was performed using the DESeq2 R package (v1.20.0) for pairwise comparisons.

### Analyze S-Sulfhydration in recombinant TFIIB by LC-MS/MS

Recombinant protein was reduced with 1 mM DTT for 30 min at 4 °C. Reactions were then passed through Zeba™ Spin Desalting Column equilibrated with PBS, pH 7.4. DTT-free proteins were then treated with hydrogen peroxide (H_2_O_2_, 5 eq. Sigma-Aldrich, cat. no. 216763) at 37 _℃_ for 30 min. Subsequently, 0.1 mM of NaHS was added for 1hr at 37 _℃_. Then 1mM IAA was added and incubated at RT for 1 h in the dark. The sample was fourfold diluted with 50 mM NH_4_HCO_3_ and digested with trypsin at a 1:50 enzyme: substrate ratio at 37 °C for 4 h. After digestion, the sample was centrifuged at 12,000 *g* for 10 min and the supernatants were dried by vacuum centrifugation. Dried samples were resuspended in 0.1% formic acid (FA) and then desalted using C18 stage tips. Digested peptides were then eluted with 50% acetonitrile and 0.1% FA. Eluted peptides were dried by vacuum centrifugation, resuspended in 0.1% FA and subjected to LC–MS/MS analysis.

### Circular dichroism spectroscopy

The secondary structures of the TFIIB were determined using far-UV circular dichroism (CD) spectra recorded with a Jasco J810 CD spectrometer (Jasco Inc., Madrid, Spain) equipped with a Jasco PTC-423S/15 Peltier accessory. The effect of NaHS on the secondary structures of TFIIB was determined by far-UV CD spectroscopy scanning from 190 to 240 nm at 37 °C. CD spectra of 1.5 mg/mL protein in 1xPBS buffer (pH 7.4) were recorded in a 0.1-cm path-length cuvette. All spectra recorded were the average of three accumulations, corrected by subtracting a buffer spectrum at 20 °C.

## Result

### Sulfur stimulates SY113 growth and drives H_2_S production

Strain SY113 was isolated from a hydrothermal vent in the Longqi hydrothermal field of the Southwest Indian Ocean at a depth of 2770 meters. The vent features a cylindrical, sulfide-rich “black smoker” chimney structure[21]. To investigate the effect of sulfur on the growth of strain SY113, we first compared its growth under conditions with and without elemental sulfur. The addition of sulfur significantly promoted the growth of SY113, a phenomenon consistent with the behavior of other known Thermococcales strains (**Fig.1a**)[13, 24]. During the stationary phase, the biomass of sulfur supplemented cultures was approximately three times greater than that of cultures without sulfur (**Fig.1b**). This indicates that while sulfur is not strictly required for growth, it substantially enhances cell proliferation. We further measured the production of hydrogen sulfide (H_2_S) under both conditions. The results showed that the H_2_S concentration in sulfur-free cultures was 17.9 ± 2.6 μM, whereas in sulfur-supplemented cultures, it increased to 170 ± 41.1 μM, an approximately tenfold rise (**Fig.1c**). This demonstrates that the externally supplied sulfur was effectively reduced to H_2_S. In summary, the addition of sulfur not only markedly promoted the growth of SY113 but was also accompanied by substantial accumulation of H_2_S, suggesting that H_2_S is a central metabolite in sulfur cycling and may be intrinsically linked to archaeal proliferation. Meanwhile, under sulfur stimulation, the transcription levels of the gene cluster encoding the membrane-bound hydrogenase MBH1, which is related to H_2_S metabolism, were significantly upregulated (**Fig. 1d**, **Fig. 1e, Supplementary Data 1**). Our recent research has shown that MBH contributes to the adaptation of strain SY113 to high-pressure conditions, suggesting that this regulatory mechanism may represent an adaptive strategy of Thermococcus species to the unique environment of deep-sea hydrothermal vent[25].

**Figure 1.**
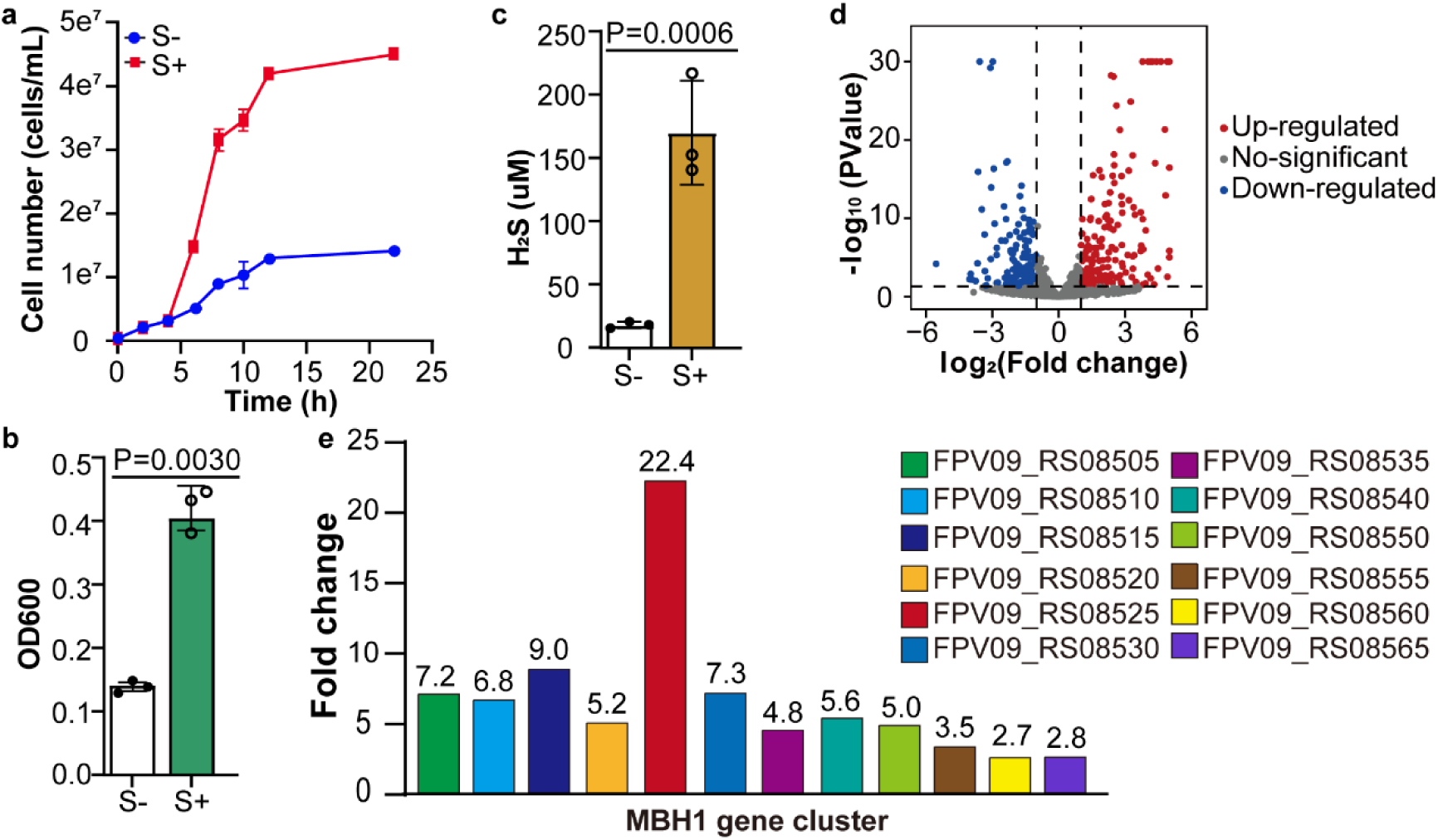
Sulfur stimulates SY113 growth and drives H2S production. **a,** Growth curves of SY113 cultivated with or without elemental sulfur, monitored by measuring the optical density at 600 nm (OD₆₀₀) over time. **b,** Biomass comparison of Sy113 with or without elemental sulfur at the stationary phase. **c,** H_2_S production by SY113 in the presence or absence of elemental sulfur. All assays were performed in triplicate. **d,** Volcano plot of transcriptomic data. The log2(sulfur/control) ratio is plotted against the -log₁₀(p-value) from a two-sample t-test. **e,** Bar chart showing the fold change in transcript levels of the MBH1 gene cluster under sulfur-replete versus sulfur-deplete conditions.

### Global reactivity profiling of protein persulfidation in SY113

Given our previous findings in mammalian/Arabidopsis cells on the unique responsiveness of the persulfidation and its involvement in functional regulatory features, we will next systematically analyze the intrinsic responsiveness of this modification in strain SY113, using a strategy similar to the one we previously described[26]. In principle, hyperreactive persulfides would saturate labeling at the low probe concentration, whereas less labile persulfides would show concentration-dependent increases in labeling. In this study, we first globally profiled persulfidation reactivity in archaea by applying Low-pH QTRP to SY113. Specifically, SY113 were lysed under pH 4.5 buffer, and then labeled with either 100 µM or 10 µM of the IPM probe. After tryptic digestion, the resulting IPM modified peptides were further conjugated to light and heavy Az-UV-biotin, respectively, via CuAAC. Then, the light and heavy labeled samples were mixed, captured with streptavidin, and photoreleased for liquid chromatography coupled with tandem mass spectrometry (LC-MS/MS) analysis (**Fig.2a**). Hyperreactive persulfides are expected to label to completion at low IPM concentrations (10 µM), and less reactive persulfides should show concentration-dependent increases in IPM labeling. Accordingly, the persulfides quantified in this analysis could be classified into three groups. Those with calculated ratio of heavy to light (100 µM vs 10 µM, R_10:1_) lower than 2.0 were defined to be hyperreactive; those within the range from 2.0 to 5.0 were considered to be moderately reactive, whereas the rest were classified as weakly reactive.

**Figure 2.**
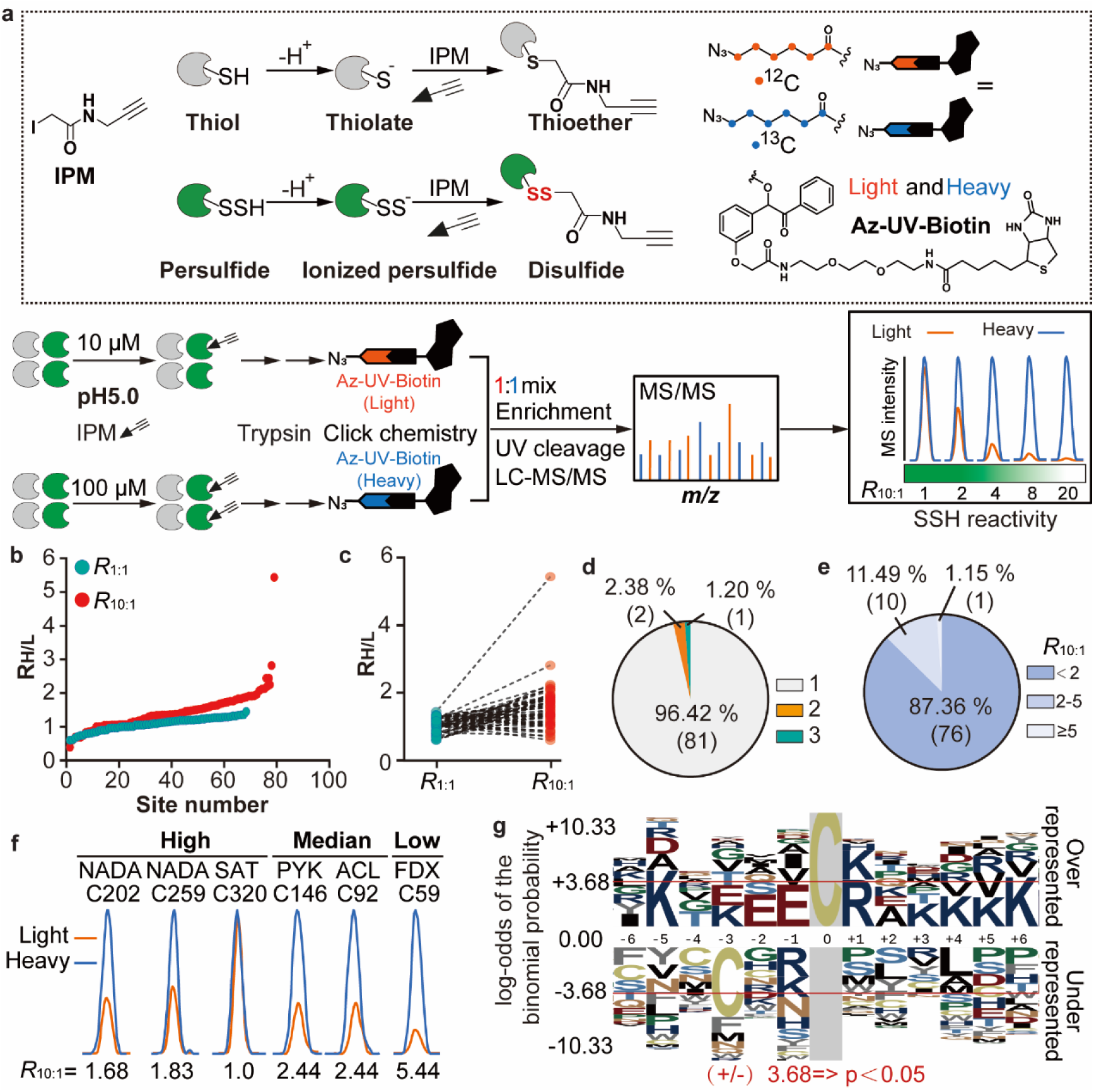
Global reactivity profiling of protein persulfidation in SY113. **a,** Workflow for analyzing the reactivity of protein persulfides. SY113 lysates were treated with the chemoselective probe IPM (10 or 100 µM). The probe-labeled proteomes were digested with trypsin, and the resulting peptides were conjugated to light or heavy Az-UV-biotin via CuAAC. The light- and heavy-labeled samples were combined at a 1:1 ratio, desalted using strong cation exchange (SCX) chromatography, and enriched with streptavidin beads. The captured peptides were then photoreleased and analyzed by LC-MS/MS. This figure outlines: the chemical structures of the IPM probe and the isotopically labeled, UV-cleavable azido-biotin, and the schematic reactions of the electrophilic IPM probe with protein persulfides or thiols. **b,** Ranking plots displaying the distribution of R_H/L_ values for R_1:1_ and R_10:1_. **c,** Side-by-side comparison of persulfides levels on the same cysteine. R_1:1_ and R_10:1_ are colored green and red, respectively. **d,** Pie chart showing the the number and percentage distribution of IPM-labeled persulfide sites. **e,** Pie charts presenting the reactivity distribution of persulfide across the SY113 proteome. **f,** Representative XICs showing the light- (yellow) and heavy- (blue) labeled IPM-peptide profiles. The mean RH/L values from biological duplicates are indicated below each XIC. **g,** Sequence motif analysis of SSH sites. Images were generated with plogo and scaled to the height of the largest column within the sequence visualization. The red horizontal lines on the plogo plots denote *p*=0.05 thresholds.

To minimize quantification bias, control experiments were performed in which lysates were treated with the same IPM concentration (i.e., 100 uM vs 100 uM, R_1:1_). High-confidence, quantifiable persulfidation sites needed to be detected in both the 10:1 and 1:1 dataset, with the latter having R values ranging from 0.67 to 1.5, with a mean value of 1.09. In total, we identified and quantified 87 -SSH sites, with a mean R_10:1_ value of 1.5, respectively (**Fig.2b, 2c, Supplementary Data 2**). In general, proteins identified in this study have a single ‘reactive’ persulfides among several quantified sites within the same polypeptide (**Fig.2d**). Compared to eukaryotes, the persulfides identified in SY113 generally exhibited higher reactivity[26]. Specifically, the vast majority (87.36%) of -SSH sites (R_10:1_ < 2) were classified as highly reactive (**Fig.2e**). Representative examples include C202 and C259 of quinolinate synthase A (NadA), C320 of sulfate adenylyltransferase (Sat), C613 of elongation factor 2 (fusA), and C158 of the TATA-box-binding protein (Tbp) (**Fig.2f**). Additionally, 11.49% of the sites showed moderate reactivity (R_10:1_ = 2-5), such as C146 of pyruvate kinase (Pyk), C92 of acetate-CoA ligase (HCL), and C374 of histidine-tRNA ligase (HTL). Only a very small proportion of sites (1.15%) displayed low reactivity (R_10:1_ > 5), exemplified by C59 of the 4Fe-4S dicluster domain-containing protein (FDX) (**Fig.2e, Fig.2f**). Notably, the C46 site of Peroxiredoxin showed extremely high reactivity (R_10:1_ = 0.39) (**Fig.S1a, b**). Phylogenetic analysis revealed that this site, which serves as a key catalytic cysteine, is evolutionarily conserved across archaea, bacteria, and eukaryotes (**Fig.S1c, d**), suggesting it may perform an ancient and fundamental biological function. To further investigate the sequence features associated with high-reactivity persulfidation, we analyzed the flanking amino acid composition of these sites. The results showed significant enrichment of charged amino acids (lysine, arginine, and aspartic acid) and alanine at positions adjacent to highly reactive sites (**Fig.2g**). Although basic and hydrophobic amino acids are also commonly enriched in the flanking regions of persulfidation sites in eukaryotes, the pronounced enrichment of an acidic amino acid (E) appears to be a unique signature of persulfidation in SY113[26, 27]. This may reflect a specific adaptation of archaeal protein regulatory mechanisms to extreme environments.

### Quantifying dynamic changes in persulfidation upon sulfur perturbation

Given the significant increase in intracellular H_2_S concentration upon sulfur addition, we hypothesized a concomitant increase in global persulfidation. To test this, we applied Low-pH QTRP to quantify the dynamic changes of persulfidation after perturbation with sulfur until growth stabilization period, then lysed and labelled with IPM. IPM-tagged proteomes with and without sulfur were digested into tryptic peptides, conjugated with light and heavy Az-UV-biotin reagents, respectively, and processed as described above. In this workflow, the heavy to light ratio calculated for each IPM-labelled persulfides provided a measure of its relative level in sulfur-treated samples versus control samples (**Fig.3a**).

In total, we identified and quantified 204 IPM-labelled persulfide sites on 171 proteins. Of these, 65.2% quantified sites showed ≥1.5-fold dynamic changes after sulfur addition (**Fig.3b, Fig.3c**). We next characterized the site-specific distribution of persulfidation in SY113 (**Fig.3d**). Among all identified proteins, the vast majority (84.8%, n = 145) contained only one cysteine subject to dynamic persulfidation regulation. These included key functional residues such as C302 of GUAB and the active-site C141 of GAP. Approximately 11.11% (19 proteins) possessed two sulfur-regulated modification sites, as exemplified by C55 and C59 in GltA, while only 4.09% (7 proteins) exhibited three or more dynamically regulated sites, with SFD1 (C138, C176, and C185) representing a notable case (**Fig. 3d, 3e**). These findings collectively demonstrate that persulfidation in SY113 is highly site-specific.

**Figure 3.**
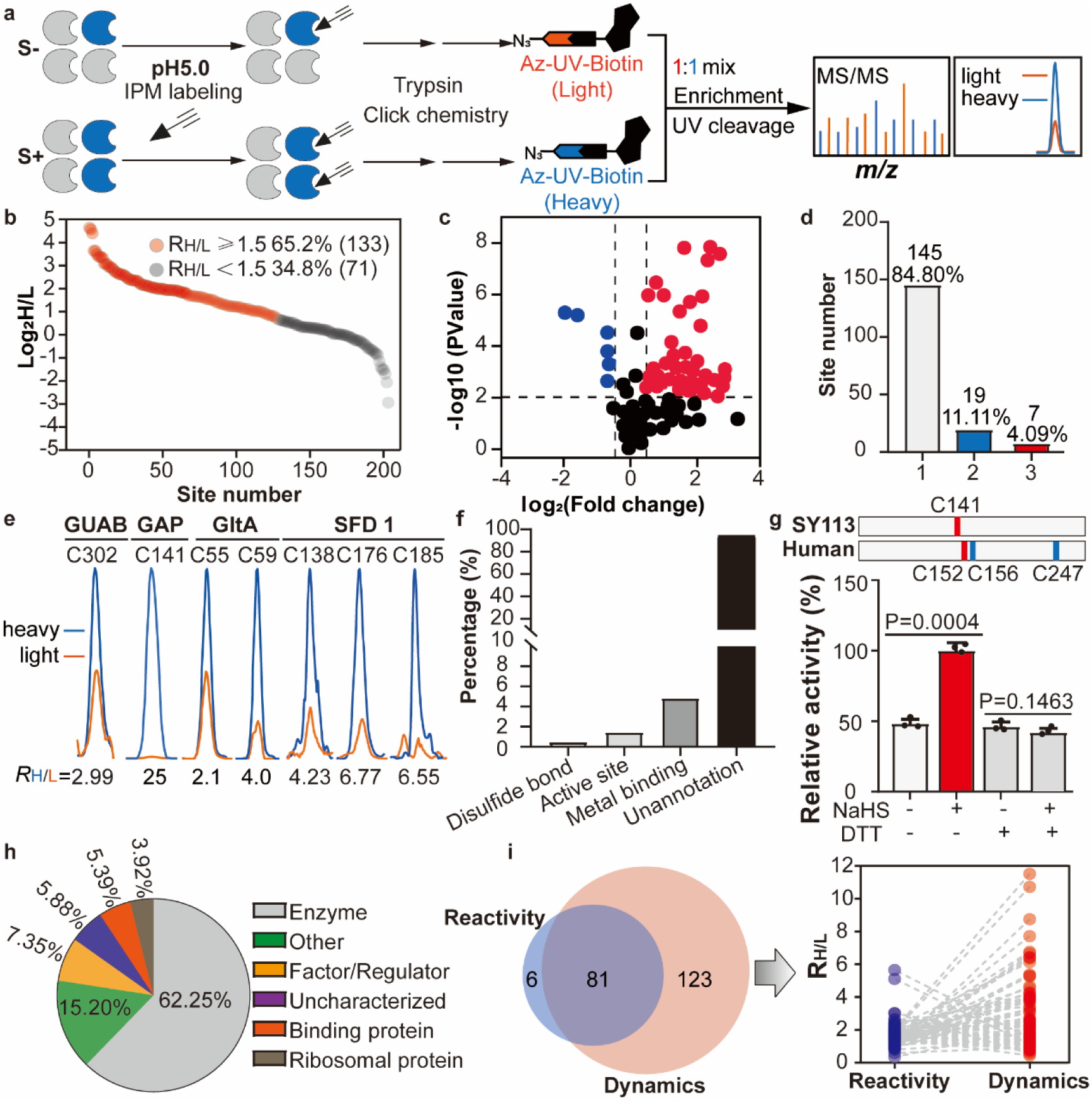
Quantifying dynamic changes in persulfidation upon sulfur perturbation. **a,** Workflow for analyzing the dynamics of persulfidation in SY113. **b,** Site-specific changes in persulfidation in response to sulfur supplementation. **c,** Volcano plot depicting the persulfidation dynamics. The log2 fold-change (sulfur-heavy/control-light) is plotted against the -log₁₀(p-value) from a two-sample t-test for all quantified cysteines. Significantly altered persulfidation events (fold change ≥ 1.5, p < 0.01) are highlighted in red. **d,** Bar chart showing the distribution of the number of probe-labeled SSH sites per protein. Number and percentage of SSH sites were shown upon each bar. **e,** Representative extracted ion chromatograms (XICs) of IPM-labeled peptides. Profiles for light- and heavy-labeled peptides are shown in yellow and blue, respectively. Mean RH/L values from biological duplicates are displayed below. **f,** Functional categorization of proteins containing persulfide sites based on UniProt annotations. **g,** Bar charts showing that the persulfidation can increase the GAP activities. Data are mean ± s.d. (representative data from biological triplicates are shown, two-sided Student’s t-test, P < 0.05 was considered significant). Each experiment was repeated three times with similar results. **h,** Pie chart showing the distribution of functional categories retrieved from the UniProt knowledge database. **i,** Venn diagram showing overlap between the reactivity and dynamics. Overlap between the reactivity and dynamics datasets. Ratios from common sites are plotted in a line series, with reactivity and dynamics shown in green and red, respectively.

Functional annotation revealed that persulfide cysteines are frequently located at structurally or functionally critical positions, including disulfide-forming residues, catalytic centers (e.g., Gap_C141), and metal-binding motifs (e.g., PriL_C364, Tfb_C26, Tdh_C111, Alas_C715, GuaB_C302, Rps14_C24, Rpl37ae_C41, and Rpl37ae_C56) (**Fig. 3f, Supplementary Data 3**). Notably, unlike most eukaryotic GAPDH enzymes that harbor three functional cysteines, the GAPDH homolog in SY113 contains only one conserved cysteine at position 141 (**Fig. 3g**). C141 exhibited pronounced persulfidation upon sulfur treatment (**Fig.3e**). Moreover, NaHS treatment markedly increased enzymatic activity, and subsequent DTT reduction returned activity, indicating that persulfidation of C141 enhances GAPDH activity, which is similar in eukaryotic organisms (**Fig. 3g**). Since GAP catalyzes the oxidation and phosphorylation of glyceraldehyde-3-phosphate to 1,3-bisphosphoglycerate with concurrent NAD+ reduction, we propose that this reversible modification may modulate glycolytic flux and cellular energy allocation under extreme environmental conditions. Furthermore, approximately 90% of the persulfide sites identified here remain functionally unannotated. Importantly, proteins harboring these poorly characterized modification sites participate in a broad spectrum of essential cellular processes, including ribosomal biogenesis, catalysis, ligase/kinase activities, metal binding, transcriptional regulation, and signal transduction (**Fig. 3h**), implying a potentially widespread role for persulfidation in modulating core physiological pathways in Thermococcales.

Building on the previously established reactivity landscape of persulfide, we investigated its relationship with dynamic regulatory behavior. We identified 81 sites common to both reactive and dynamically modified pools, accounting for 93.1% of all reactive sites and 39.7% of dynamically regulated sites (**Fig. 3i**). Statistical evaluation revealed a positive correlation between site reactivity and susceptibility to sulfur-mediated regulation. In particular, low-reactivity sites (e.g., FeSD_C59, reactivity = 5.44, dynamism = 1.3) were largely unaffected by sulfur regulation. A subset of medium-reactivity sites, such as PYK_C146 (reactivity = 2.44, dynamism = 1.73), showed regulatory responses, whereas 63.8% of high-reactivity sites displayed clear sulfur-dependent regulation, as seen in TBP_C158 and TFB_C26. Although exceptions existed, for instance, PXPA_C26 (reactivity = 1.19) was not regulated, the data strongly support the notion that high-reactivity cysteines are more prone to sulfur-mediated dynamic modification (**Fig. S2**).

### Persulfidation modulates diverse cellular processes in SY113

Upon in-depth analysis of the aforementioned data, we detected persulfidation modification of cysteine residues within the redox-active CXXC motif. The CXXC motif typically consists of a peroxidatic cysteine (C(P)) and a resolving cysteine (C(R)) [28, 29]. Among the identified persulfidation sites, a total of 16 sites were located at C(P), including FDX_C48, SAT_C320, and TFIIB_C26, among others, while 4 sites were located at C(R), such as RPS14_C24 and DevR_C81 (**Fig. 4a**, **Fig. 4b**). Proteins harboring these persulfidation were primarily classified as enzymes, transcription factors, binding proteins, and ribosomal proteins (**Fig. 4c**). Notably, C26, the C(P) site within the CXXC motif of the general transcription factor TFIIB, is regulated by hydrogen sulfide (**Fig. 4b**). TFIIB is a key component in the assembly of the transcription initiation complex in both eukaryotes and archaea[30–32]. The C26 participates in forming a tetrahedral coordination structure, binding Zn²_⁺_ and thereby maintaining the normal structure and function of this zinc-binding protein (**Fig. 4d**). After *in vitro* expression and purification of TFIIB, NaHS stimulation followed by mass spectrometry analysis confirmed that the C26 undergoes persulfidation (**Fig. 4e**). Circular dichroism spectroscopy further demonstrated that NaHS treatment induced conformational changes in the TFIIB (**Fig. 4f**). In addition, sulfate adenylyltransferase (SAT) catalyzes the first step of the sub-pathway from sulfate to sulfite, which is part of the H_2_S biosynthetic pathway and is further integrated into the sulfur metabolic network[25, 33]. C320 is located within the ATP-sulfurylase domain of SAT, suggesting its potential involvement in the enzyme’s catalytic function. Through structural prediction using AlphaFold and comparison with the known structure 1V47 from *Thermus thermophilus* HB8[34], we found C320, along with C332 and C323, forms a metal-binding pocket that coordinates a Zn²_⁺_ ion, thereby stabilizing the protein’s spatial conformation (**Fig. 4g**). Further *in vitro* experiments showed that NaHS treatment significantly enhanced SAT enzyme activity, while subsequent reduction with DTT restored its activity to baseline levels (**Fig. 4h**). These results confirm that persulfidation at SAT_C320 has an activating effect, indicating that this modification may regulate SAT’s conformation and activity by influencing Zn²_⁺_ binding capacity, thereby participating in the regulation of cellular sulfur metabolism. In summary, this study demonstrates that persulfidation within the CXXC motif can broadly regulate the conformation and biological functions of the affected proteins, providing new molecular insights into the role of H_2_S in protein post-translational modifications and related metabolic regulatory networks.

**Figure 4.**
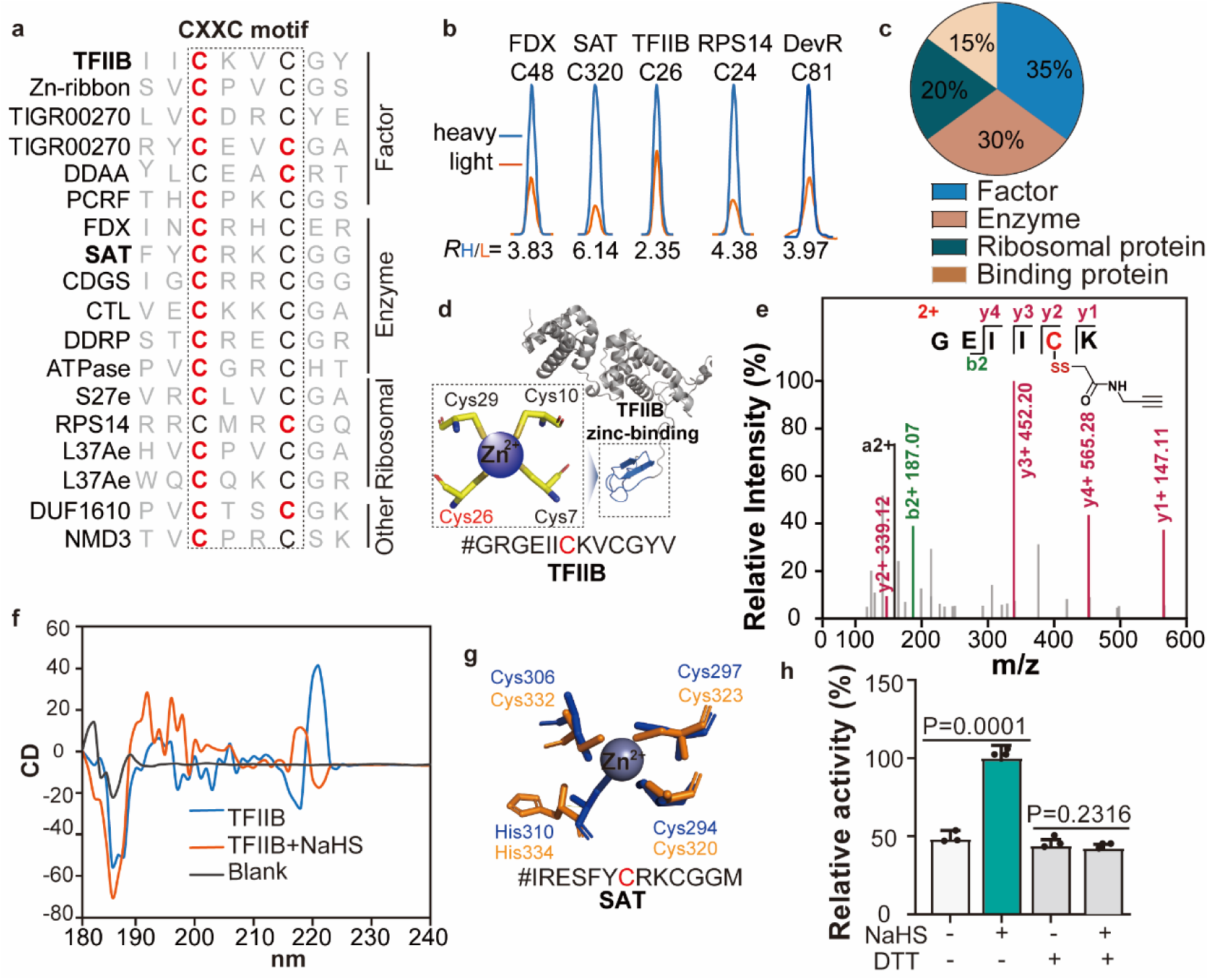
Persulfidation modulates diverse cellular processes in SY113. **a,** Sequence showing CXXC motif amino acid sequences. The persulfide sites are highlighted in red. **b,** Representative extracted ion chromatograms (XICs) of IPM-labeled peptides. Light- and heavy-labeled peptide profiles are shown in red and blue, respectively. Mean RH/L values from biological duplicates are indicated below each XIC. **c,** Pie chart showing functional categorization of CXXC motif-containing proteins based on UniProt annotations. **d,** Cartoon representation of the TFIIB protein structure (AlphaFold model) generated in PyMOL 2.1.1. The metal-binding pocket is displayed in stick representation. **e,** MS/MS spectrum unambiguously identifying Cys26 of TFIIB as a persulfidation site via the IPM-tagged peptide. **f,** Circular dichroism spectroscopy confirms that persulfidation induces conformational changes in TFIIB. **g,** Structural superposition of the SAT protein from *Sy113* (AlphaFold model, yellow) and the ortholog from *Thermus thermophilus* HB8 (PDB: 3IVZ, blue). Structures are shown in stick representation and were aligned using PyMOL 2.1.1. **h,** Bar charts showing persulfidation enhances SAT enzymatic activity. Data are mean ± s.d. from one representative experiment performed in triplicate. Statistical significance was determined by a two-sided Student’s t-test (*p < 0.05). The experiment was independently repeated three times with consistent results.

## Discussion

This study employed a chemical proteomics to systematically unveil, for the first time, the widespread presence and biological significance of protein persulfidation in the deep-sea hyperthermophilic archaeon *T.aciditolerans* SY113. Our work not only expands the field of H_2_S signaling biology to the third domain of life but, more importantly, suggests that this ancient post-translational modification may play a pivotal role in enabling organisms to adapt to extreme environments.

The most fundamental finding of this research is that persulfidation in SY113 is not a stochastic event but exhibits high site selectivity, unique reactivity features, and significant dynamic regulation (**Fig.2**, **Fig.3**). The marked enrichment of acidic amino acids flanking the modification sites, which is distinct from patterns observed in eukaryotes, may reflect unique evolutionary adaptations in the archaeal protein regulatory machinery to high-temperature and acidic conditions[26, 27]. Furthermore, the significant positive correlation we discovered between highly reactive sites and their sulfur powder-induced dynamic changes strongly suggests these modifications represent precisely regulated biological processes rather than non-specific chemical byproducts. Our functional validation of GAPDH (Gap_C141) provides direct evidence: persulfidation directly activates this enzyme’s activity. This indicates that H_2_S can rapidly adjust central energy metabolic flux by modifying key metabolic enzymes, potentially a crucial strategy for SY113 to cope with fluctuations in environmental energy sources[35]. Particularly striking is our discovery of extensive persulfidation within the ancient redox-sensing module, the CXXC motif. This finding directly links H_2_S signaling to an ancient regulatory mechanism[13]. In-depth investigation of TFIIB and SAT preliminarily outlines a complete regulatory circuit from metabolism to transcription: H_2_S, through modification of the general transcription factor TFIIB, directly intervenes in the transcription initiation machinery, ultimately coordinating the organism’s adaptation to environmental stresses such as high pressure[22]. Concurrently, environmental sulfur is metabolized to produce H_2_S, which then, by modifying SAT (a key enzyme in sulfur metabolism), may establish a fine-tuned feedback loop. This proposes a new paradigm: in the sulfur-rich environment of deep-sea hydrothermal vents, persulfidation acts as a central bridge connecting environmental sulfur signals, intracellular H_2_S levels, and the regulation of overall physiological function. It represents a core molecular strategy for *Thermococcus* to adapt to their unique ecological niche.

Despite the groundbreaking progress, this study has several limitations that also chart directions for future research. Firstly, the functional validation for the majority of the identified modification sites remains at the level of inference and correlation. Although we have demonstrated the functional impact on key proteins like GAPDH, SAT, and TFIIB through *in vitro* experiments, direct evidence for the physiological consequences of persulfidation at specific sites within the archaeal cell has not yet been obtained. Future work should involve constructing site-directed mutant strains (e.g., mutating cysteine to serine) to directly assess the contribution of this modification to protein function, signaling pathways, and ultimately, the overall fitness of SY113 *in vivo*. Secondly, our research focused on a single strain and a single environmental perturbation. The dynamics of persulfidation across different growth phases and in response to other environmental pressures (such as temperature, pH, or high hydrostatic pressure) remain to be elucidated. Future time-series and multi-factorial perturbation analyses will more precisely delineate the role of this modification within the stress response network. Finally, the crosstalk between persulfidation and other redox modifications (such as sulfenylation and S-nitrosylation), and how this modification network integrates into the broader metabolic and gene expression regulatory networks within the unique domain of Archaea, remains to be explored[36]. Deciphering this ancient regulatory network is crucial not only for understanding extremophile adaptation but will also provide valuable molecular clues for exploring the origins of signal transduction mechanisms during early life evolution.

In summary, this study opens new avenues for H_2_S signal transduction research in Archaea and establishes persulfidation as an ancient and important regulatory mechanism. Future research will continue to delve into the molecular details of this modification network and its profound implications for the evolution of life.

## Structural illustration

The structure of the SAT, GAP and TFIIB was downloaded from uniport, which were from AlphaFold, the structure of SAT from *T. thermophilus* HB8 was downloaded from PDB: 1V47. All the protein structure graphics were generated with PyMOL Molecular Graphics System (https://pymol.org/2/).

## Statistical analysis

Statistical analysis was performed using Excel (Microsoft 2021) or Prism 9 (GraphPad) software. The number of biological replicates and statistical methods are indicated in the figure legends. Statistical significance was calculated using the two-sided unpaired Student’s t-test or two-way ANOVA. All error bars show standard deviations (SD) of the mean. Statistical significance was defined as p values ≤ 0.05.

## Data availability

The main data supporting the findings of this study are available within the article and its Supplementary Information files. Additional details on datasets and protocols that support the findings of this study will be made available by the corresponding author upon reasonable request. The raw data of the proteomics generated in this study have been deposited in iProX database under accession code PXD07017. and are publicly available. Source data are provided with this paper.

## Acknowledgements

This work was supported by grants from the National Key R&D Program of China (2022YFA1303000) and the National Natural Science Foundation of China (31800036, 42376123, 32270126)

## Author contributions

L.F. performed the chemoproteomics and functional experiments, analyzed the data, and wrote the manuscript, L.X.G prepare SY113 strains for chemoproteomics and transcriptome analysis. S.W.L and Z.Y.H purified the protein, performed a TFIIB persulfidation analysis, and conducted circular dichroism spectroscopy analysis, J.Y., L.F. and L.X.G conceived the project, supervised the work, analyzed data and the wrote the manuscript with inputs from others.

---

**Supplementary Figure 1.**
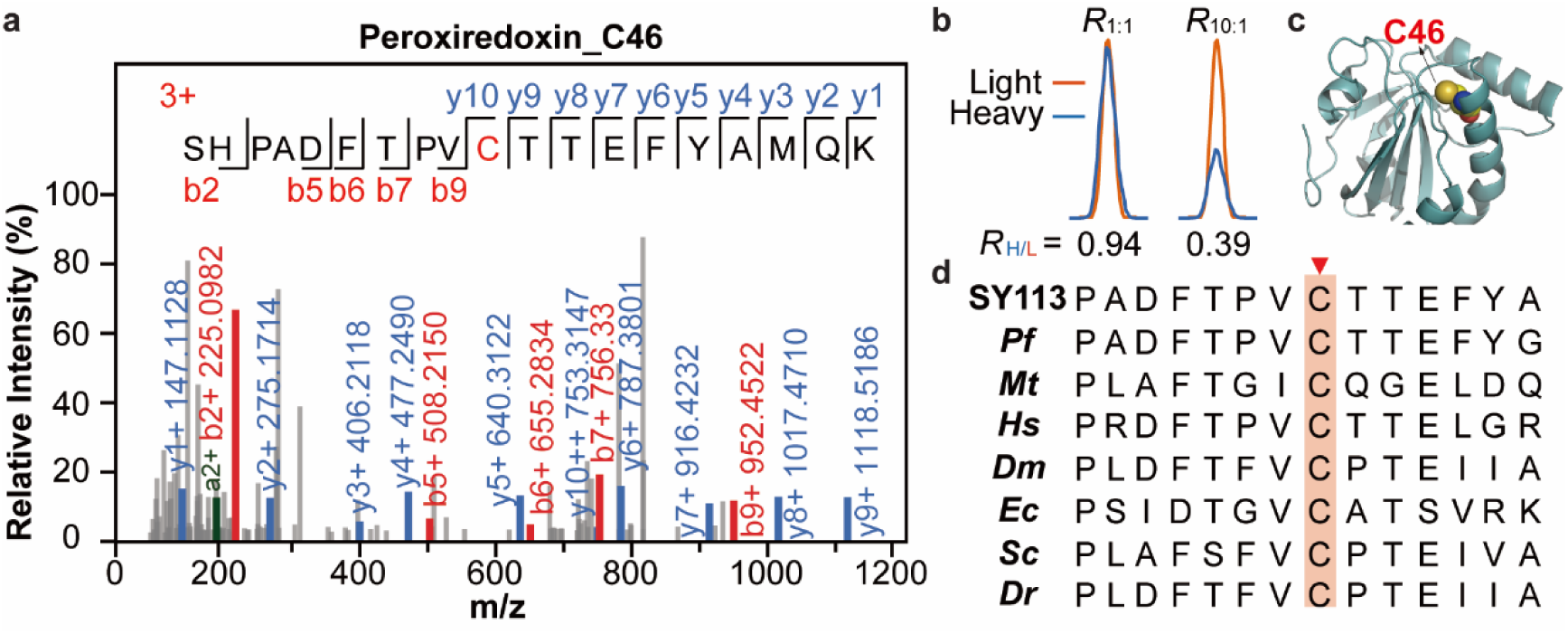
Characterization of Peroxiredoxin persulfidation. **a,** Representative MS ion chromatograms for the Peroxiredoxin C46 peptide as a target site of persulfidation. **b**, Representative XICs showing changes in IPM-labeled peptides from Peroxiredoxin. The profiles for light and heavy-labeled peptide are shown in yellow and blue, respectively. The average *R*_H/L_ values calculated from biological duplicates are displayed below each XIC. **c**, 3D protein structure of Peroxiredoxin were visualized with Pymol 2.1.1. **d**, Multiple sequence alignment showing peroxiredoxin amino acid sequences. The highly conserved active site is underlined. Representatives shown from Eukaryota (Dm=*Drosophila melanogaster*; Hs=*Homo sapiens*; Sc=*Saccharomyces cerevisiae*; Dr=*Danio rerio*), Bacteria (Ec=*Escherichia coli*; Mt=*Mycobacterium tuberculosis*) and Archaea(SY113, Pf=*Pyrococcus furiosus*). Critical residues in the active site of peroxiredoxins (in red and highlight in red triangle) are conserved in all organisms.

**Supplementary Figure 2.**
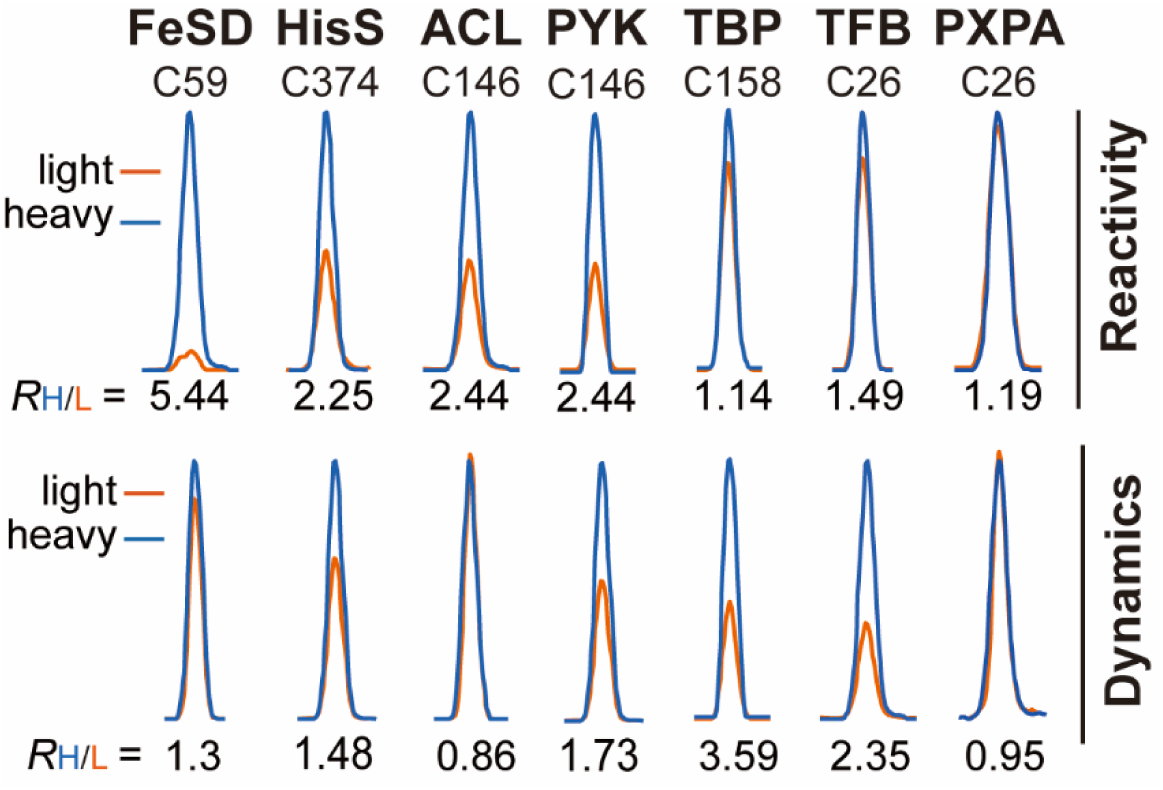
Correlation of persulfidation reactivity and dynamics. Representative XICs showing changes in IPM-labeled peptides from reactivity (up) and dynamics (down). The profiles for light and heavy-labeled peptide are shown in yellow and blue, respectively. The average *R*_H/L_ values calculated from biological duplicates are displayed below each XIC.

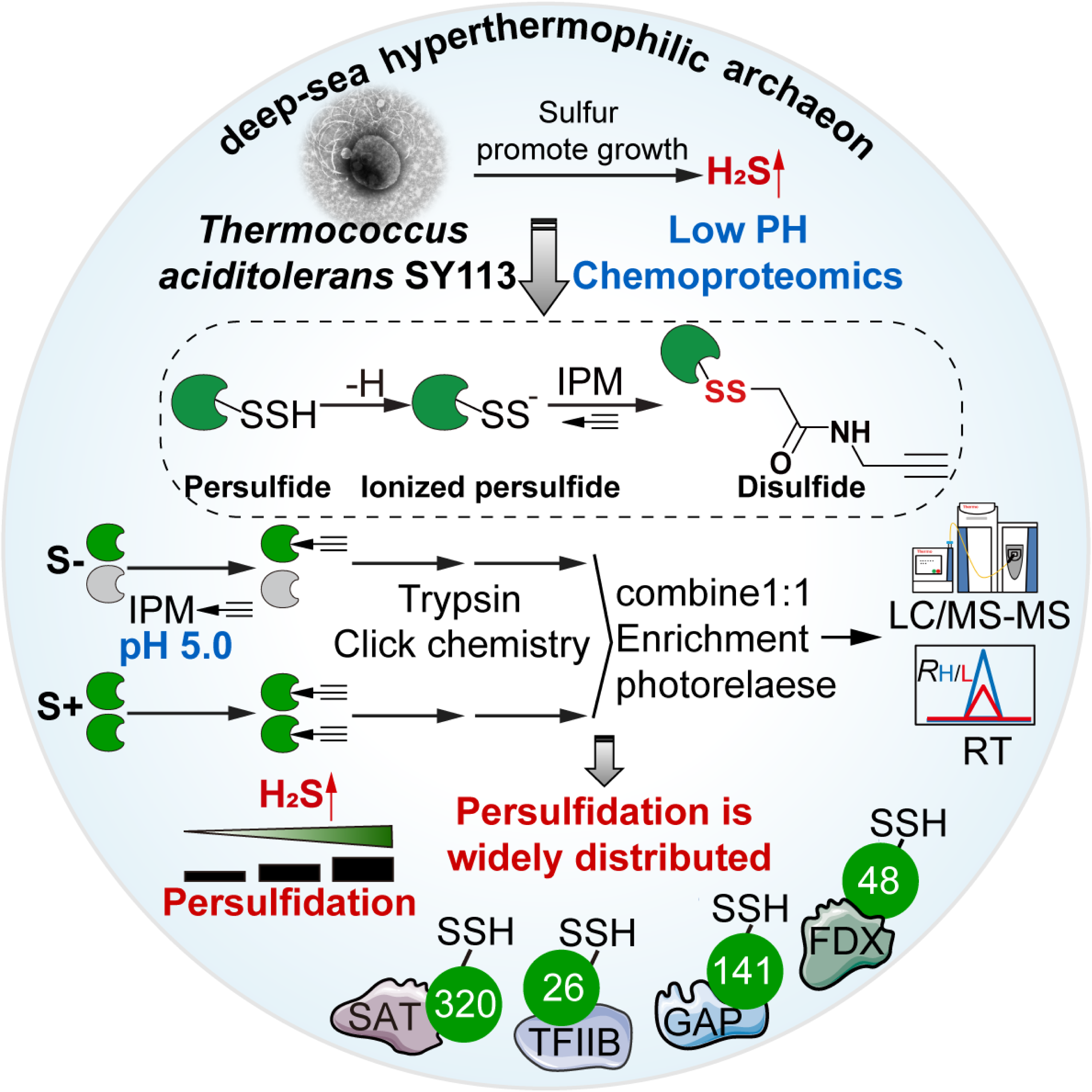

